# DrugPTM-Bench: A Large-Scale Dataset for Predictive Modeling of Drug-Induced Cell Type-Specific Protein Post-Translational Modifications

**DOI:** 10.64898/2026.04.27.721113

**Authors:** Amitesh Badkul, Mohammadsadeq Mottaqi, Li Xie, Lei Xie

## Abstract

Protein post-translational modifications (PTMs), particularly phosphorylation, serve as the primary “molecular switches” that orchestrate cellular signaling and drug response. While PTM dysregulation is a hallmark of cancer and neurodegeneration, the lack of standardized, drug-perturbed datasets has hindered the development of predictive models capable of capturing context-dependent PTM responses. Effective predictive modeling must therefore integrate multidimensional data, including the specific drug, dosage, treatment duration, cellular background, and the modified site. However, existing PTM resources remain largely static and fail to capture drug-induced regulation across these critical dimensions. To address this gap, we present DrugPTM-Bench, a curated, large-scale benchmark derived from decryptM-derived dose-dependent PTM measurements, standardizing site-level drug response across 7 cancer cell lines, 27 drugs, and 11,167 proteins. Comprising 99.5% phosphorylation events, the dataset includes six time points, 16 dosage levels, and pEC50 potency values (half-maximal effective concentration). We formulate a classification task to identify upregulated, downregulated, or unchanged PTM sites (following a drug treatment), a critical step in deciphering drug Mechanism of Action (MoA) and target engagement. Our evaluation reveals that in protein-disjoint out-of-distribution (OOD) setting, baseline machine learning and deep learning models struggle to recover minority regulation classes, while standard rebalancing strategies improve recall only at the cost of precision and overall F1-score. These results indicate that current methods do not learn robust decision boundaries between regulated and unchanged PTM events. DrugPTM-Bench provides a phosphoproteomics benchmark for modeling drug-induced PTM regulation in imbalanced biological settings. Beyond classification, DrugPTM-Bench’s retention of pEC50 values, drug perturbation profiles, and site-level sequence context enables additional predictive tasks including drug potency regression, mechanism-of-action prediction from PTM fingerprints, and drug-specific PTM site sensitivity ranking, establishing a multi-task benchmark for PTM-centric drug discovery. Ultimately, DrugPTM-Bench establishes a rigorous framework for developing robust, context-aware models to elucidate drug MoA and signaling dynamics.

## Introduction

Protein post-translational modifications (PTMs) are biochemical processes that regulate protein function, stability, localization, and interactions through chemical alterations. These modifications typically involve the addition of functional groups such as acetyl, methyl, or phosphoryl to specific amino acids (39; 53; 45). PTMs regulate essential biological functions, including cell differentiation, metabolism, and immune response. They also influence signal transduction, gene expression, DNA repair, and the cell cycle. By influencing protein conformation, structure, lifespan, activity, charge, solubility, and interactions with DNA, RNA, and other proteins, PTMs ultimately shape cellular function and phenotypic outcomes (35; 53). These modifications occur at different stages of biological processes across multiple cellular compartments, including the Golgi apparatus, endoplasmic reticulum, nucleus, and cytoplasm (5).

Beyond their fundamental biological roles, PTMs also modulate signaling pathways, create novel druggable sites, and enhance the specificity of therapeutic interventions (40). Dysregulation of PTMs is implicated in numerous diseases, particularly cancer, neurological disorders, and metabolic syndromes, making them crucial therapeutic targets. For example, phosphorylation dysregulation is a hallmark of cancer, leading to the targeting of kinases in drug development (15). Inhibiting overactive kinases can prevent the phosphorylation of key proteins involved in uncontrolled cell growth and survival (18). Similarly, targeting ubiquitination pathways, such as proteasome inhibitors, has led to successful treatments for multiple myeloma (22). Aberrant PTMs can also contribute to protein misfolding and aggregation, as seen in neurodegenerative disorders like Alzheimer’s and Parkinson’s, where modifying PTM pathways presents a promising therapeutic approach (1). Drug-induced PTM changes are pharmacologically informative because they provide dynamic readouts of pathway perturbation that can help illuminate target engagement and mechanism of action (52). Despite their significance, only about 7% of biological systems of PTMs have been explored using small molecule ligands, highlighting the vast, uncharted chemical space for PTM-targeted drug development (40).

To better characterize PTMs, researchers rely on experimental techniques such as mass spectrometry (MS) coupled with immunoaffinity enrichment, which enables precise identification and quantification of protein modifications (20). Another widely used approach, Proximity Ligation Assay (PLA), employs antibody-based proximity detection combined with DNA amplification for signal enhancement (45). However, both methods have significant drawbacks. MS is expensive and requires extensive sample preparation, while PLA depends on antibody specificity and availability, limiting its applicability (28). The ability to systematically characterize PTMs is critical for understanding cellular regulation, drug response, and disease pathology. However, experimental PTM profiling remains costly and technically demanding, which has motivated the development of computational approaches for PTM prediction and analysis. In contrast to experimental methods, computational PTM prediction provides a fast and low-cost strategy for proteome-wide annotation and experimental design, and machine learning has become an option for PTM-site prediction (45; 49; 51). While machine learning (ML) and deep learning (DL) have significantly improved PTM prediction and analysis (34; 2; 10; 50), existing PTM databases including dbPTM, PhosphoSitePlus, and BioGRID, primarily capture static PTM annotations rather than dynamic, drug-perturbed PTM responses (11; 21; 44). As a result, they are limited in their ability to support modeling of how PTM states change across drug, dose, time, and cellular context. Recent efforts, such as decryptM (51), have addressed this gap by systematically profiling dose- and time-resolved PTM responses across multiple drugs and cell lines, generating over 1.8 million dose-response curves. Importantly, decryptM shows that drug-regulated PTMs can shed light on target engagement and drug mechanism of action, underscoring the pharmacological relevance of dynamic PTM measurements (51). While decryptM provides comprehensive, quantitative PTM profiling across drugs, doses, cell lines, and time points, it is not organized as a benchmark for predictive modeling: it lacks standardized tasks, curated splits, and explicit labels for site-level regulatory outcomes required for machine learning evaluation. To bridge this gap, we introduce DrugPTM-Bench, a curated benchmark that transforms dose-dependent PTM (ddPTM) measurements from the decryptM resource into standardized, site-level prediction tasks enriched with explicit experimental context. DrugPTM-Bench captures PTM regulation across cell lines, drugs, dosage levels, and time points, and also retains pEC50 values, enabling both classification and regression-oriented formulations. In the present work, we focus primarily on PTM regulation classification: given a PTM site, a drug, and its treatment condition, the task is to predict whether the site is upregulated, downregulated, or unchanged following treatment. This formulation is pharmacologically meaningful because drug-induced PTM changes are direct readouts of perturbed signaling pathways and therefore relevant to understanding drug response (51; 52; 49).

Another contribution of DrugPTM-Bench is that it exposes a biologically realistic imbalanced-learning regime. Across PTM-chemical pairs, the majority of sites exhibit neutral regulation, creating substantial class imbalance when predicting regulatory outcomes. In some benchmark settings, the neutral-to-minority ratio reaches at least 165:1 (Fig. 2).

This matters computationally because imbalanced data are known to bias classifiers toward majority classes, while common resampling strategies can introduce noise or degrade decision boundaries. Most work on imbalance in machine learning addresses this problem through resampling or augmentation strategies, ranging from classical methods such as the Synthetic Minority Oversampling Technique (SMOTE) (7) and the Adaptive Synthetic Sampling Approach (ADASYN) (19) to more recent contrastive-learning-based approaches (9; 25). However, these methods are typically developed and evaluated in domains where the data structure differs substantially from biological perturbation data, and synthetic resampling can be especially problematic when minority examples are sparse and biologically heterogeneous. Existing imbalance benchmarks such as CIFAR-100-LT (32) and ImageNet-LT (38), are also rooted in computer vision and do not capture the context dependence or label structure of PTM regulation data. Consistent with this pattern, the baseline ML and DL models in our benchmark struggle to recover minority regulation classes, and rebalancing improves recall only at the expense of precision and overall F1-score. DrugPTM-Bench therefore serves first as a benchmark for dynamic drug-induced PTM regulation modeling, and second as a benchmark for learning under imbalanced, context-dependent biological data In summary, our main contributions are: (1) We introduce DrugPTM-Bench, a large-scale, standardized benchmark dataset for drug-induced, context-dependent PTM regulation, constructed from decryptM-derived ddPTM measurements; (2) We formally define and benchmark the site-level PTM regulation prediction task, providing standardized splits and baseline ML/DL results under extreme class imbalance; (3) We empirically demonstrate and analyze the limitations of existing imbalance-handling methods in this biological context; (4) Lastly, we identify and motivate three additional predictive tasks enabled by DrugPTM-Bench beyond the primary classification task, including pEC50 potency regression, MoA prediction from PTM perturbation profiles, and drug-specific PTM site sensitivity ranking, establishing a roadmap for PTM-centric computational drug discovery (see Appendix A).

## Results and Discussion

### Dataset

#### Drug-induced PTM regulation dataset and benchmark Existing Dataset Gap

Existing PTM datasets have primarily evolved along three axes: (1) expansion of curated PTM sites, (2) functional and mechanistic annotation, and (3) integration with biological networks. Early resources, such as Phospho.ELM (12), focused on cataloging phosphorylation sites, providing conservation scores and disorder metrics but lacking drug-disease associations. PhosphoSitePlus (21) expanded PTM annotations to over 450,000 sites, linking them to disease mutations, yet it retained a static view of PTMs without capturing dose- and time-dependent drug effects. EPSD 2.0 (8) integrated functional annotations for 88,074 phosphorylation events, incorporating multi-omics data, yet its drug-PTM associations were inferred rather than experimentally validated. dbPTM (11) further integrated 2.8 million PTM sites with kinase-substrate and E3 ligase interactions, allowing hypothesis generation about upstream regulators, but without providing explicit drug perturbation effects over time. Similarly, ActiveDriverDB (30) maps disease-associated mutations to PTM regions but lacks information on how pharmacological interventions modulate PTMs. As summarized in Table 1, no existing dataset systematically captures dose- and time-resolved drug-induced PTM regulations, highlighting the need for DrugPTM-Bench as a dedicated resource for studying dynamic PTM perturbations in response to drug treatments.

**Table 1.**
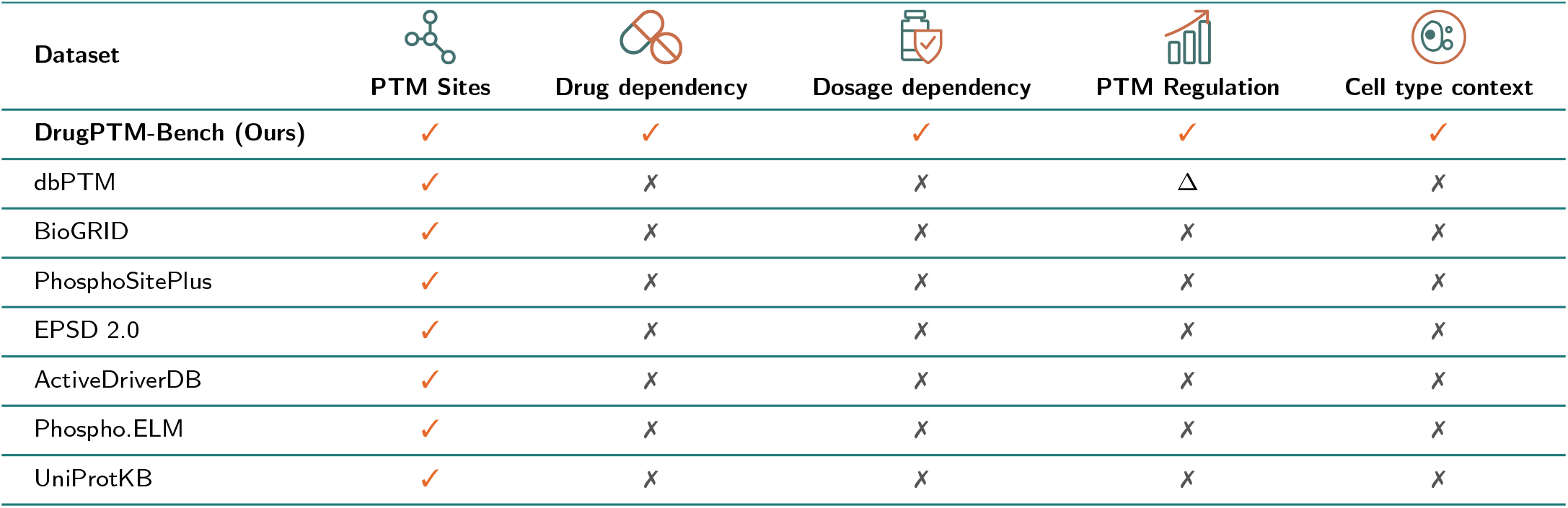
Comparison of different PTM datasets with DrugPTM-Bench. (Δ denotes partial PTM regulation info).

| Dataset | PTM Sites | Drug dependency | Dosage dependency | PTM Regulation | Cell type context |
| --- | --- | --- | --- | --- | --- |
| **DrugPTM-Bench (Ours)** | ✓ | ✓ | ✓ | ✓ | ✓ |
| dbPTM | ✓ | ✗ | ✗ | $\Delta$ | ✗ |
| BioGRID | ✓ | ✗ | ✗ | ✗ | ✗ |
| PhosphoSitePlus | ✓ | ✗ | ✗ | ✗ | ✗ |
| EPSD 2.0 | ✓ | ✗ | ✗ | ✗ | ✗ |
| ActiveDriverDB | ✓ | ✗ | ✗ | ✗ | ✗ |
| Phospho.ELM | ✓ | ✗ | ✗ | ✗ | ✗ |
| UniProtKB | ✓ | ✗ | ✗ | ✗ | ✗ |

#### Dataset Overview

DrugPTM-Bench is constructed from decryptM-derived ddPTM (dose-dependent post-translational modification) measurements and comprises 10,733,521 PTM records across seven human cancer cell lines, 27 unique drugs, 11,167 proteins, six treatment time points, and 16 dosage levels. In addition to PTM identities and protein sequence context, the benchmark retains the key experimental variables required for predictive modeling, including cell line, drug, dosage, treatment time, signal ratio, and negative logarithmic half-maximal effective concentration (pEC50) values. Fig. 1 provides a high-level overview of the benchmark composition and response distributions.

**Figure 1.**
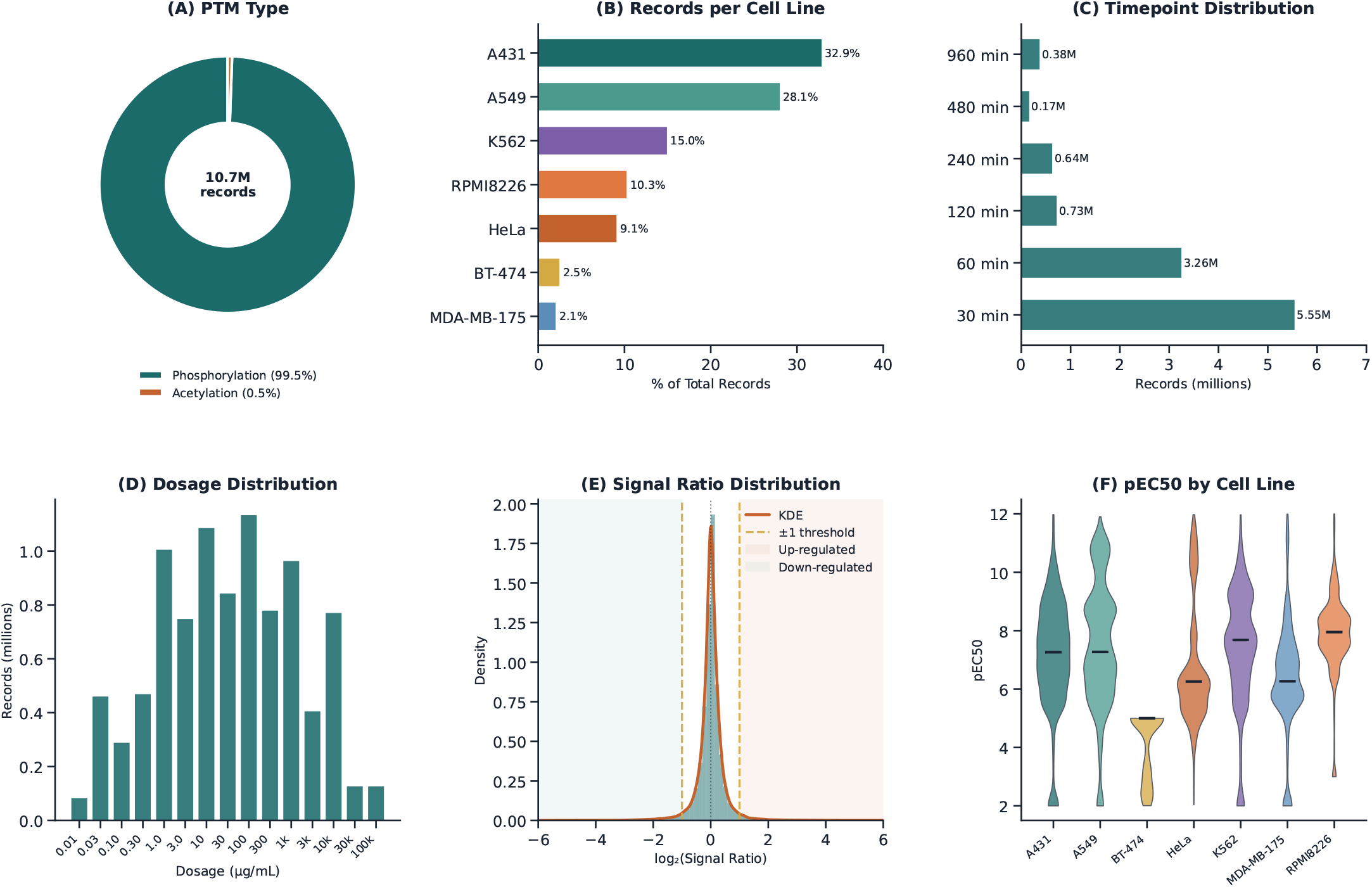
DrugPTM-Bench dataset overview. Six-panel summary of the dataset. (A) PTM type distribution: 99.5% phosphorylation, 0.5% acetylation. (B) Number of PTM measurements per cell line. (C) Timepoint distribution (30–960 min). (D) Dosage level distribution (0.01 – 100,000 µg/mL, 16 levels). (E) log2 Signal Ratio distribution across all cell lines, showing the characteristic spike at zero (neutral sites). (F) pEC50 distribution by cell line

As shown in Fig. 1A, DrugPTM-Bench is dominated by phosphorylation measurements (99.5%), while acetylation accounts for only a small minority of records (0.5%). Given this strong skew in PTM type composition, the current benchmark experiments focus primarily on phosphorylation-based regulation modeling. This unevenness reflects the underlying experimental design of the source perturbation proteomics data rather than any additional balancing imposed during benchmark construction. The benchmark captures PTM responses over six post-treatment time points and 16 dosage levels spanning several orders of magnitude. As shown in Fig. 1C, measurements are concentrated at earlier response windows, particularly 30 and 60 minutes, indicating that the dataset is enriched for acute drug-induced PTM responses.

Fig. 1D shows that dosage coverage is broad, ranging from 0.01 to 100,000 *µ*g/mL, with the highest record density concentrated in the intermediate dosage range. Together, these two additional dimensions make DrugPTM-Bench suitable for studying how PTM regulation varies across both treatment duration and drug exposure level. To quantify PTM response magnitude, DrugPTM-Bench uses the signal ratio between treated and untreated conditions, which is subsequently analyzed on the log_2_ scale. Fig. 1E shows that the log_2_(signal ratio) distribution is sharply centered near zero, indicating that most PTM sites remain close to unchanged under a given treatment condition, while a smaller subset exhibits strong positive or negative shifts. This overall distribution already suggests that regulated PTM events are relatively sparse compared with neutral responses, a property that becomes central in the benchmark task formulation described later. Finally, DrugPTM-Bench retains pEC50 values as a measure of drug potency at the PTM-site level. As shown in Fig. 1F, pEC50 values span a broad range across cell lines, reflecting substantial variation in the strength and sensitivity of PTM responses under drug perturbation.

#### Signal ratio, regulation labels, and benchmark tasks

To quantify drug-induced PTM response, DrugPTM-Bench uses the signal ratio (SR), defined as the ratio of PTM signal intensity in the drug-treated condition to that in the untreated control condition:

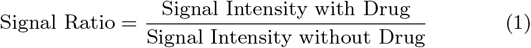

This quantity provides a direct measure of the magnitude and direction of drug-induced PTM change at a site. Because PTM abundances span a wide dynamic range, we analyze SR on the log_2_ scale. The log_2_ transformation is standard in proteomics because it compresses large fold-change ranges and renders up- and down-regulation symmetric around zero, thereby making the response more suitable for statistical analysis and machine learning.

Fig. 1E shows that the log_2_(SR) distribution is sharply centered near zero, with most PTM sites exhibiting little or no response under a given treatment condition. A closer examination of the interval between −3 and 3 shows that 136,925 PTM-drug pairs (1.2% of all pairs) have log_2_(SR) > 1, whereas 168,538 pairs (1.5%) have log_2_(SR) < −1. Following a widely used proteomics convention, we treat | log_2_(SR)| > 1 as a biologically meaningful response threshold. Accordingly, each PTM measurement is assigned one of three regulation labels: *Up* for log_2_(SR) > 1, *Down* for log_2_(SR) < −1, and *Neutral* for −1 ≤ log_2_(SR) ≤ 1. This converts the quantitative PTM response into an explicit three-class prediction problem.

Constructing a sequence-resolved benchmark further requires mapping each PTM-containing peptide to a protein sequence. This is non-trivial in bottom-up proteomics because peptides can be shared among multiple proteins, making protein inference inherently ambiguous. In the decryptM source data, MaxQuant (48) reports peptide groups together with one or more candidate proteins, of which a leading protein is selected as the most representative entry for downstream analysis. DrugPTM-Bench handles this ambiguity through a leading-protein-first mapping strategy: PTM peptides are first mapped to the MaxQuant leading protein, and only when no valid match is found is the search extended to additional candidate proteins. This procedure does not eliminate peptide-origin ambiguity entirely, but it yields a consistent and biologically grounded mapping for sequence-based modeling. Using this strategy, we obtain explicit PTM-site positions on full-length protein sequences, which are required for constructing model-ready benchmark instances. DrugPTM-Bench supports two related predictive formulations. Among the mapped entries, 25 proteins had more than one leading protein assignment, and 21 proteins could not be retained because their identifiers had been deleted from ENSEMBL (14), leaving 10,827 leading proteins in the final benchmark. The positions of all mapped PTM sites are included in DrugPTM-Bench.

#### Class imbalance landscape

A central property of DrugPTM-Bench is the severe class imbalance inherent to drug-induced PTM regulation. As shown in Fig. 2B, the vast majority of PTM measurements remain neutral across all cell lines, with only a small fraction of sites classified as up- or down-regulated. However, this imbalance is not uniform across cellular contexts. In the aggregate view across all dosage conditions, A549 exhibits the strongest overall imbalance, with only 1.1% of measurements falling into the regulated classes, whereas RPMI8226 shows the weakest overall imbalance, with 8.3% regulated sites. K562 is also strongly imbalanced in the aggregate view, with only 1.5% regulated measurements. Fig. 2A further shows that imbalance depends strongly on dosage. The most extreme ratios occur at lower dosage bins, where regulated PTM events are especially sparse. For example, cell line K562 reaches a neutral-to-regulated ratio of 165:1 in the lowest dosage bin, and A549 reaches 131:1 in the 0.01–0.1 *µ*g/mL range. As dosage increases, the imbalance generally decreases across most cell lines, indicating that stronger drug exposure perturbs a larger fraction of PTM sites. This pattern is consistent with dose-resolved phosphoproteomic studies showing that increasing drug concentration enhances target and pathway engagement and propagates perturbations more broadly through signaling networks (51). Nevertheless, even at the highest dosage bins, the neutral class remains dominant in nearly all cell lines.

**Figure 2.**
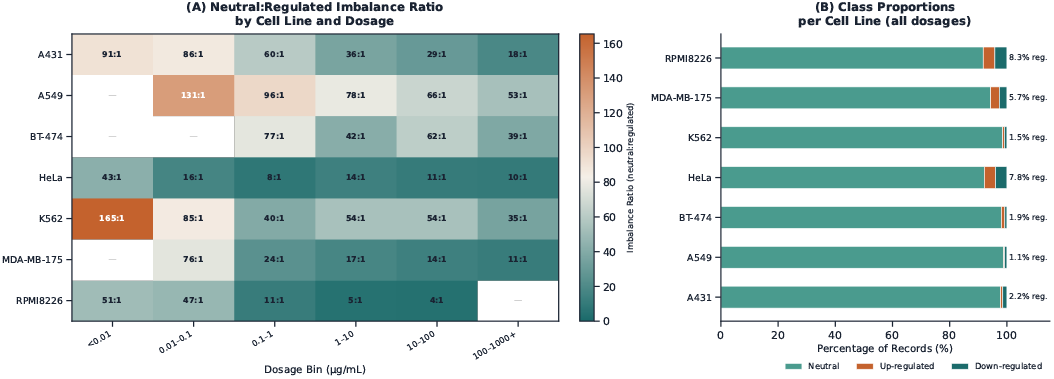
Class imbalance landscape across cell lines. (A) Heatmap of neutral-to-minority class ratios across cell lines and dosage levels. (B) Stacked bar chart showing the proportion of Up, Down, and Neutral PTM sites per cell line. Imbalance ratios range from 4:1 (RPMI8226) to 165:1 (K562).

A biologically plausible interpretation is that the imbalance landscape reflects the breadth of pathway perturbation induced by the underlying drug panel in each cellular background. Drug classes with broad downstream effects, such as proteasome inhibitors, are known to induce widespread signaling and proteostatic changes, whereas more selective perturbations may leave most PTM sites unchanged (16). This interpretation is consistent with the relatively low imbalance observed in RPMI8226 and the more severe imbalance in cell lines where the measured perturbations appear to affect a narrower fraction of the PTM landscape. For the purposes of benchmarking, the key implication is that DrugPTM-Bench does not present a single fixed imbalance ratio. Instead, it captures a spectrum of imbalance regimes across cell lines and dosage conditions, providing a more realistic setting for evaluating context-dependent PTM regulation models.

#### Drug-induced perturbation landscape

Fig. 3 summarizes how strongly different drugs perturb the PTM landscape across cellular contexts. Fig. 3A aggregates all non-neutral PTM responses, defined by the benchmark regulation labels (*Up* and *Down*), for each drug-cell line combination. Bubble size represents the total number of regulated PTM sites, whereas bubble colour reflects the balance between up-and down-regulation, allowing perturbation magnitude and directionality to be visualized simultaneously. This view shows that drug-induced PTM perturbation is highly heterogeneous across cellular contexts: some drug-cell line combinations induce broad PTM reprogramming, whereas others affect only a comparatively small subset of sites. Such heterogeneity with the dose-resolved PTM measurements can reveal target engagement, pathway engagement, and mechanism-of-action-related signatures in a context-dependent manner.

**Figure 3.**
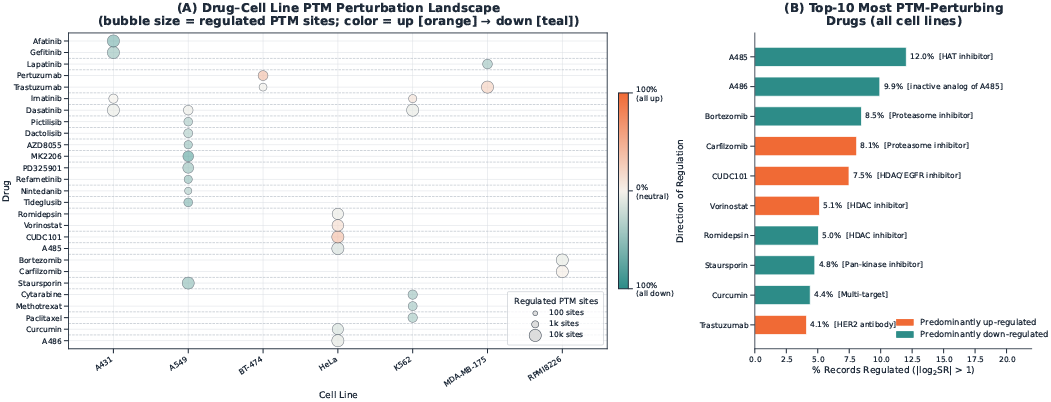
Drug-induced PTM perturbation landscape across drugs and cell lines. (A) Bubble plot summarizing regulated PTM responses for each drug-cell line combination after applying the benchmark regulation threshold (| log_2_ SR| *>* 1). Bubble size indicates the total number of regulated PTM sites (Up + Down), and bubble colour indicates the fraction of regulated sites that are up-regulated, ranging from predominantly down-regulated (teal) to predominantly up-regulated (orange). (B) Top-10 perturbing drugs ranked by the percentage of PTM records classified as regulated in the benchmark. The bracketed mechanism labels are literature-derived annotations added for interpretive context. These annotations highlight that the most strongly perturbing compounds span diverse therapeutic mechanisms, including chromatin regulation, proteasome inhibition, receptor/kinase signalling blockade, and antibody-based targeting.

**Figure 4.**
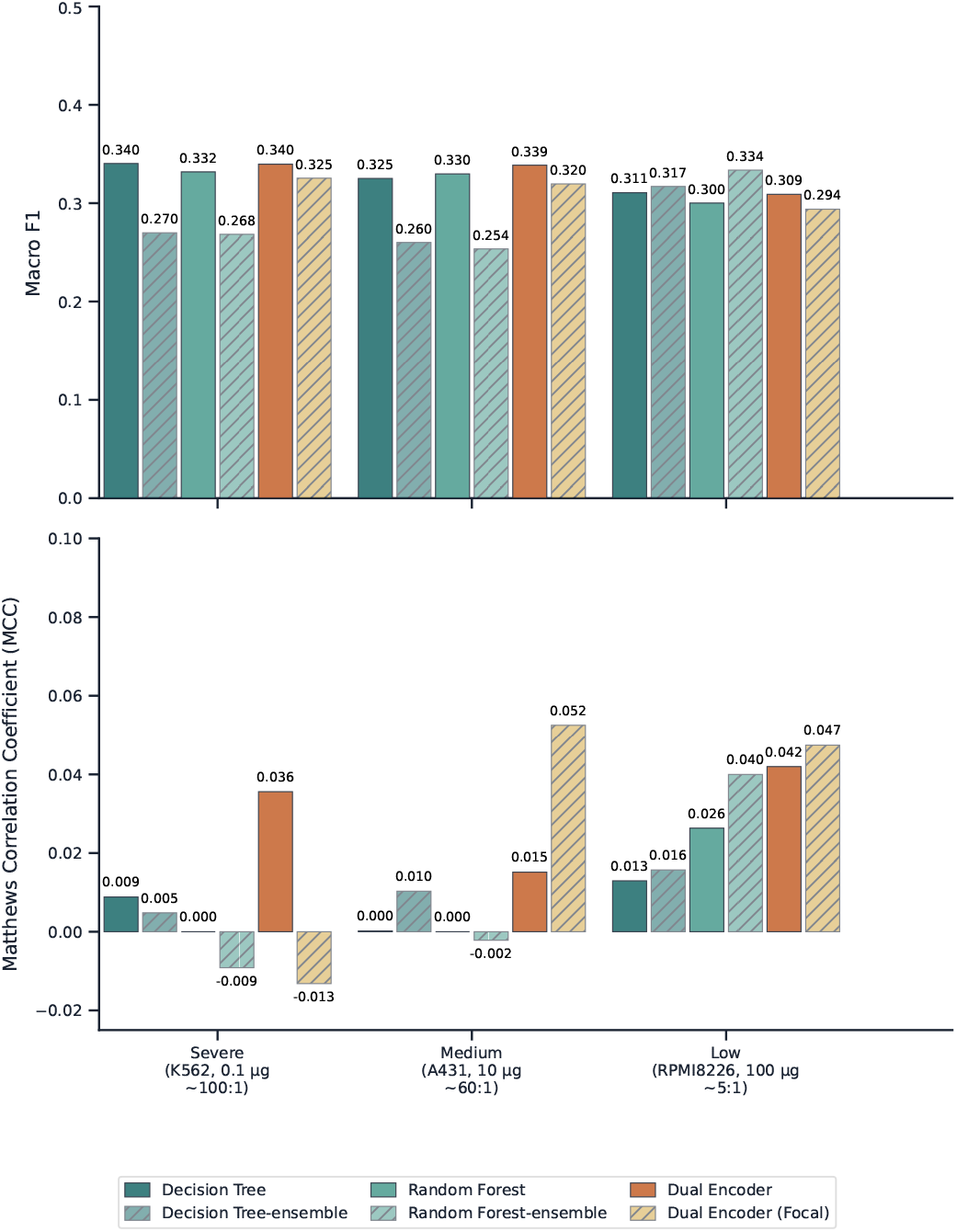
Benchmark performance across three imbalance regimes using macro-F1 and Matthews correlation coefficient (MCC). The top panel reports macro-F1 and the bottom panel reports MCC for ML and DL baselines. Macro-F1 remains relatively similar across several settings despite major differences in minority-class recoverability, whereas MCC more clearly distinguishes meaningful performance under imbalance.

Fig. 3B ranks the most perturbing drugs in the benchmark. The highest-ranking compounds are enriched for drugs with broad effects on chromatin regulation, proteostasis, and signalling, including the CBP/p300 histone acetyltransferase inhibitor A-485 (33), the proteasome inhibitors bortezomib and carfilzomib (29), the multi-target HDAC/EGFR/HER2 inhibitor CUDC-101 (6), the HDAC inhibitors vorinostat and romidepsin (17; 4), the pan-kinase inhibitor staurosporine (43), and the HER2-targeted monoclonal antibody trastuzumab (31). A-486 should not be described as an active HAT inhibitor (33), rather, it is better treated as the inactive analog or control compound associated with A-485. This spectrum of mechanisms suggests that DrugPTM-Bench contains a wide range of perturbational regimes, from chromatin-directed and proteostasis-directed responses to receptor- and kinase-driven signaling effects, thereby increasing the benchmark’s value for modeling context-dependent PTM regulation.

#### Additional Predictive Tasks Enabled by DrugPTM-Bench

The perturbation landscape described above illustrates that DrugPTM-Bench captures rich, multi-dimensional drug-PTM relationships that extend well beyond the primary classification task. Specifically, the retention of pEC50 values enables a drug potency regression task at PTM-site resolution, the availability of full drug perturbation profiles across sites enables mechanism-of-action prediction, and the site-level signal ratios per drug-protein pair enable a drug-specific PTM site sensitivity ranking task. Each of these formulations addresses a distinct open problem in computational drug discovery for which no existing benchmark currently operates at PTM-site resolution. A structured description of each task, including its input and output specification, biological motivation, gap relative to existing benchmarks, and key supporting literature is provided in Appendix A.

### Benchmark

#### Problem Description

Given a PTM site, a drug, and its dosage, the objective is to predict the regulatory outcome, whether the PTM site is upregulated, downregulated, or remains neutral following treatment. To illustrate the impact of DrugPTM-Bench’s class imbalance which ranges from low to medium to extreme, we frame this as a classification task, where models must learn to distinguish between rare regulatory events and dominant neutral cases. We evaluate the baseline models across three benchmark settings that span a spectrum of class imbalance: **Severe** (K562 cell line, 0.1 *µ*g/mL, neutral-to-minority ratio ≈100:1), **Medium** (A431 cell line, 10 *µ*g/mL, ≈60:1), and **Low** (RPMI8226 cell line, 100 *µ*g/mL, ≈5:1). The class counts for each setting are summarized in Table 2. To ensure an out-of-distribution (OOD) evaluation, a protein-disjoint split is applied: proteins are randomly partitioned into training, validation, and test sets in a 7:2:1 ratio, guaranteeing that no protein appears in more than one split. This design mimics the real-world scenario in which models must be generalized to previously unseen protein targets.

**Table 2.**
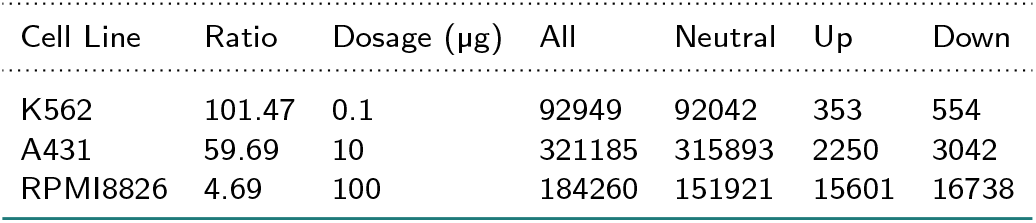
Extreme class imbalance in phosphorylation site counts for prediction of PTM regulation at various dosages in A431 and HeLa cell lines, 30 minutes after treatment.

**Table 3.**
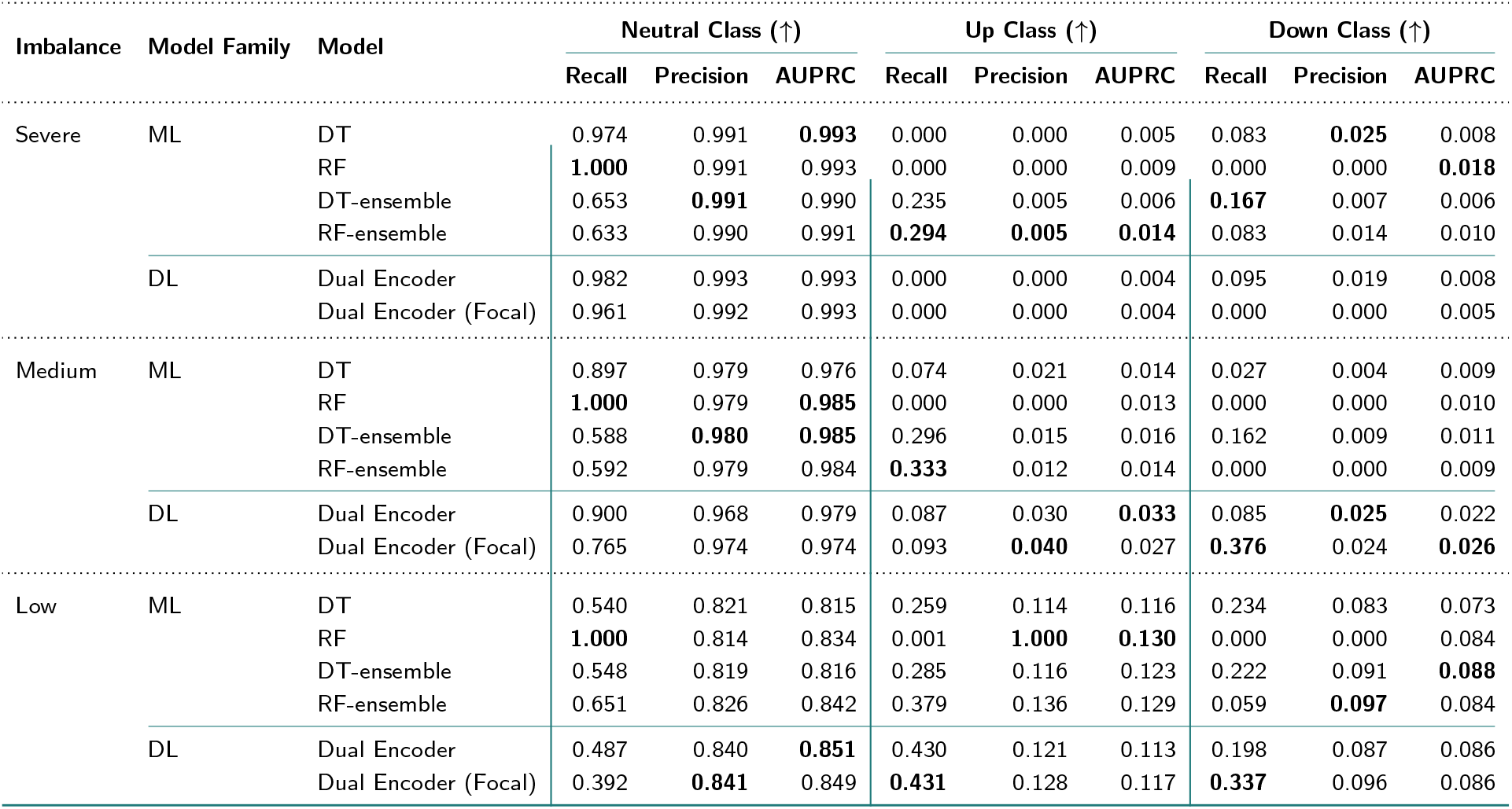
Benchmark performance across imbalance regimes. Ensemble models denote training on rebalanced subsets. Metrics are grouped by class to highlight class-specific performance. Arrows indicate that higher is better (↑).

| Imbalance | Model Family | Model | Neutral Class (↑) |  |  | Up Class (↑) |  |  | Down Class (↑) |  |  |
| --- | --- | --- | --- | --- | --- | --- | --- | --- | --- | --- | --- |
|  |  |  | Recall | Precision | AUPRC | Recall | Precision | AUPRC | Recall | Precision | AUPRC |
| Severe | ML | DT | 0.974 | 0.991 | **0.993** | 0.000 | 0.000 | 0.005 | 0.083 | **0.025** | 0.008 |
|  |  | RF | **1.000** | 0.991 | 0.993 | 0.000 | 0.000 | 0.009 | 0.000 | 0.000 | **0.018** |
|  |  | DT-ensemble | 0.653 | **0.991** | 0.990 | 0.235 | 0.005 | 0.006 | **0.167** | 0.007 | 0.006 |
|  |  | RF-ensemble | 0.633 | 0.990 | 0.991 | **0.294** | **0.005** | **0.014** | 0.083 | 0.014 | 0.010 |
|  | DL | Dual Encoder | 0.982 | 0.993 | 0.993 | 0.000 | 0.000 | 0.004 | 0.095 | 0.019 | 0.008 |
|  |  | Dual Encoder (Focal) | 0.961 | 0.992 | 0.993 | 0.000 | 0.000 | 0.004 | 0.000 | 0.000 | 0.005 |
| Medium | ML | DT | 0.897 | 0.979 | 0.976 | 0.074 | 0.021 | 0.014 | 0.027 | 0.004 | 0.009 |
|  |  | RF | **1.000** | 0.979 | **0.985** | 0.000 | 0.000 | 0.013 | 0.000 | 0.000 | 0.010 |
|  |  | DT-ensemble | 0.588 | **0.980** | **0.985** | 0.296 | 0.015 | 0.016 | 0.162 | 0.009 | 0.011 |
|  |  | RF-ensemble | 0.592 | 0.979 | 0.984 | **0.333** | 0.012 | 0.014 | 0.000 | 0.000 | 0.009 |
|  | DL | Dual Encoder | 0.900 | 0.968 | 0.979 | 0.087 | 0.030 | **0.033** | 0.085 | **0.025** | 0.022 |
|  |  | Dual Encoder (Focal) | 0.765 | 0.974 | 0.974 | 0.093 | **0.040** | 0.027 | **0.376** | 0.024 | **0.026** |
| Low | ML | DT | 0.540 | 0.821 | 0.815 | 0.259 | 0.114 | 0.116 | 0.234 | 0.083 | 0.073 |
|  |  | RF | **1.000** | 0.814 | 0.834 | 0.001 | **1.000** | **0.130** | 0.000 | 0.000 | 0.084 |
|  |  | DT-ensemble | 0.548 | 0.819 | 0.816 | 0.285 | 0.116 | 0.123 | 0.222 | 0.091 | **0.088** |
|  |  | RF-ensemble | 0.651 | 0.826 | 0.842 | 0.379 | 0.136 | 0.129 | 0.059 | **0.097** | 0.084 |
|  | DL | Dual Encoder | 0.487 | 0.840 | **0.851** | 0.430 | 0.121 | 0.113 | 0.198 | 0.087 | 0.086 |
|  |  | Dual Encoder (Focal) | 0.392 | **0.841** | 0.849 | **0.431** | 0.128 | 0.117 | **0.337** | 0.096 | 0.086 |

**Table 4.**
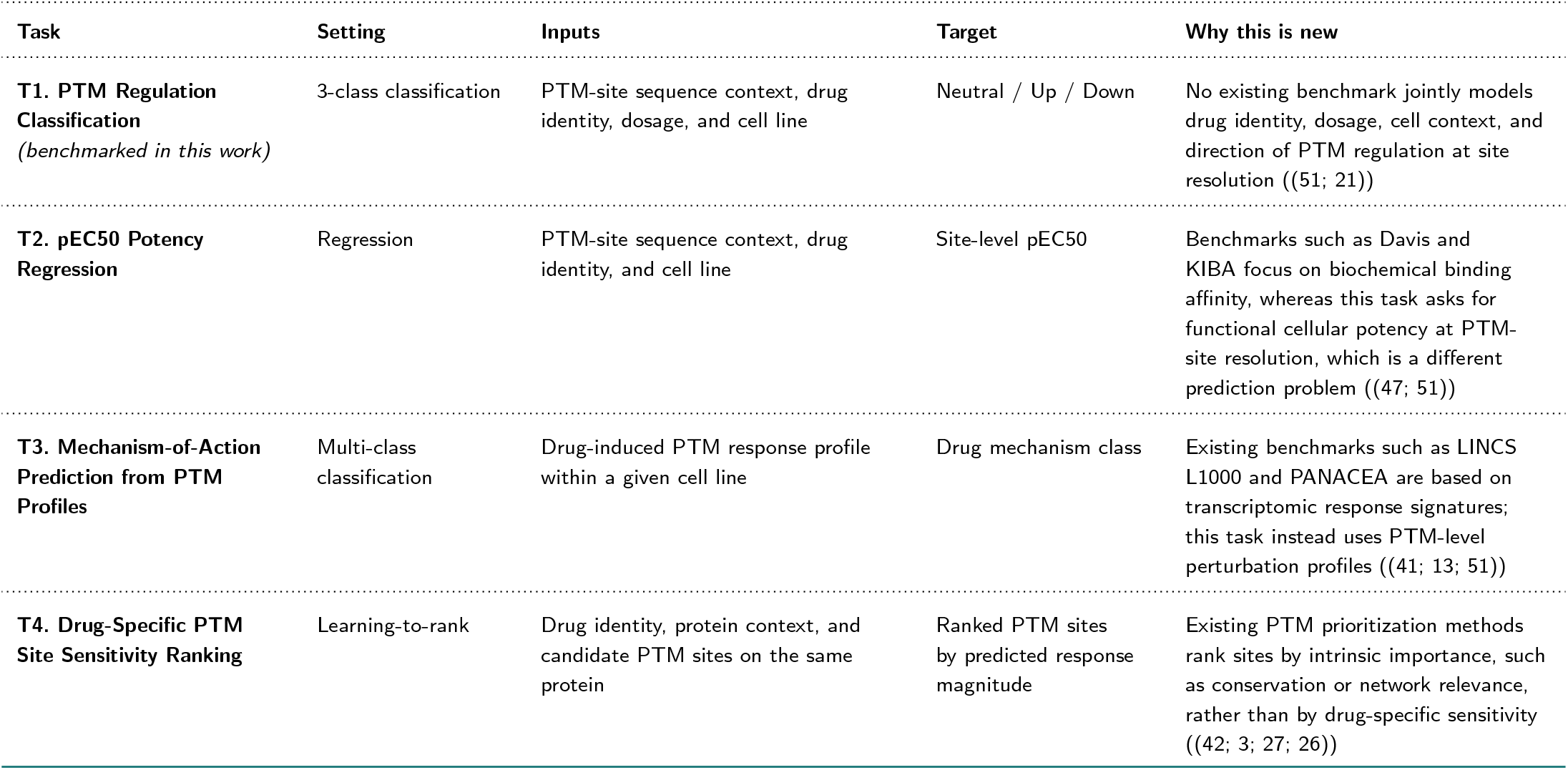
Predictive tasks enabled by DrugPTM-Bench. T1 is benchmarked in this work. T2-T4 are defined here as future benchmark tasks. All tasks are formulated at PTM-site resolution.

| Task | Setting | Inputs | Target | Why this is new |
| --- | --- | --- | --- | --- |
| **T1. PTM Regulation Classification**<br>*(benchmarked in this work)* | 3-class classification | PTM-site sequence context, drug identity, dosage, and cell line | Neutral / Up / Down | No existing benchmark jointly models drug identity, dosage, cell context, and direction of PTM regulation at site resolution ((51; 21)) |
| **T2. pEC50 Potency Regression** | Regression | PTM-site sequence context, drug identity, and cell line | Site-level pEC50 | Benchmarks such as Davis and KIBA focus on biochemical binding affinity, whereas this task asks for functional cellular potency at PTM-site resolution, which is a different prediction problem ((47; 51)) |
| **T3. Mechanism-of-Action Prediction from PTM Profiles** | Multi-class classification | Drug-induced PTM response profile within a given cell line | Drug mechanism class | Existing benchmarks such as LINCS L1000 and PANACEA are based on transcriptomic response signatures; this task instead uses PTM-level perturbation profiles ((41; 13; 51)) |
| **T4. Drug-Specific PTM Site Sensitivity Ranking** | Learning-to-rank | Drug identity, protein context, and candidate PTM sites on the same protein | Ranked PTM sites by predicted response magnitude | Existing PTM prioritization methods rank sites by intrinsic importance, such as conservation or network relevance, rather than by drug-specific sensitivity ((42; 3; 27; 26)) |

**Table 5.** Extreme class imbalance in phosphorylation site counts for prediction of PTM regulation at various dosages and time points in the benchmark dataset.

| Cell line | Dosage<br>(μg) | Timepoint<br>(min) | All | Neutral | Up | Down |
| --- | --- | --- | --- | --- | --- | --- |
| A431 | 0 | 30 | 174758 | 174758 | non | non |
| A431 | 0.03 | 30 | 174661 | 172673 | 907 | 1081 |
| A431 | 0.3 | 30 | 174646 | 172437 | 837 | 1372 |
| A431 | 1 | 30 | 174697 | 172871 | 1075 | 751 |
| A431 | 3 | 30 | 174692 | 171893 | 1385 | 1414 |
| A431 | 10 | 30 | 174683 | 171866 | 1256 | 1561 |
| A431 | 30 | 30 | 174709 | 170714 | 1456 | 2539 |
| A431 | 100 | 30 | 174684 | 169190 | 1602 | 3892 |
| A431 | 300 | 30 | 174686 | 168631 | 1727 | 4328 |
| A431 | 1000 | 30 | 174675 | 168820 | 1586 | 4269 |
| A431 | 10000 | 30 | 174686 | 165355 | 2246 | 7085 |
| A549 | 0 | 60 | 274619 | 273817 | 397 | 405 |
| A549 | 1 | 60 | 157947 | 156730 | 400 | 817 |
| A549 | 3 | 60 | 126852 | 125798 | 351 | 703 |
| A549 | 10 | 60 | 157819 | 156034 | 584 | 1201 |
| A549 | 30 | 60 | 126760 | 125102 | 519 | 1139 |
| A549 | 100 | 60 | 157900 | 155684 | 747 | 1469 |
| A549 | 300 | 60 | 126757 | 124875 | 577 | 1305 |
| A549 | 1000 | 60 | 126762 | 124683 | 586 | 1493 |
| A549 | 3000 | 60 | 126677 | 124502 | 692 | 1483 |
| A549 | 10000 | 60 | 126635 | 124152 | 799 | 1684 |
| HeLa | 0 | 30 | 20367 | 20367 | non | non |
| HeLa | 0 | 240 | 45539 | 42274 | 1541 | 1724 |
| HeLa | 0 | 960 | 12335 | 12315 | 12 | 8 |
| HeLa | 0.03 | 960 | 6144 | 6097 | 23 | 24 |
| HeLa | 0.1 | 960 | 6146 | 6090 | 21 | 35 |
| HeLa | 0.3 | 960 | 6144 | 6071 | 33 | 40 |
| HeLa | 1 | 960 | 6154 | 5822 | 96 | 236 |
| HeLa | 3 | 960 | 6148 | 5907 | 101 | 140 |
| HeLa | 10 | 30 | 20351 | 17472 | 1017 | 1862 |
| HeLa | 10 | 960 | 6150 | 5740 | 167 | 243 |
| HeLa | 30 | 30 | 20357 | 17999 | 1126 | 1232 |
| HeLa | 30 | 240 | 6247 | 6078 | 30 | 139 |
| HeLa | 30 | 960 | 6142 | 5724 | 176 | 242 |
| HeLa | 100 | 30 | 20360 | 19062 | 806 | 492 |
| HeLa | 100 | 240 | 22764 | 22193 | 293 | 278 |
| HeLa | 100 | 960 | 6141 | 5644 | 180 | 317 |
| HeLa | 300 | 30 | 20351 | 18250 | 944 | 1157 |
| HeLa | 300 | 240 | 22726 | 22015 | 341 | 370 |
| HeLa | 1000 | 30 | 20355 | 16637 | 1403 | 2315 |
| HeLa | 1000 | 240 | 22760 | 22188 | 339 | 233 |
| HeLa | 3000 | 30 | 20313 | 19765 | 194 | 354 |
| HeLa | 3000 | 240 | 22801 | 22249 | 314 | 238 |
| HeLa | 10000 | 30 | 20355 | 14766 | 2218 | 3371 |
| HeLa | 10000 | 240 | 22806 | 21873 | 461 | 472 |
| HeLa | 30000 | 30 | 20347 | 16846 | 1278 | 2223 |
| HeLa | 30000 | 240 | 22805 | 21887 | 510 | 408 |
| HeLa | 100000 | 30 | 20339 | 18876 | 634 | 829 |
| HeLa | 100000 | 240 | 22783 | 21621 | 691 | 471 |
| K562 | 0 | 30 | 83950 | 83950 | non | non |
| K562 | 0.01 | 30 | 44385 | 44237 | 103 | 45 |
| K562 | 0.03 | 30 | 68663 | 68358 | 188 | 117 |

Table 5 – continued from previous page
| Cell line | Dosage<br>($\mu\text{g}$) | Timepoint<br>(min) | All | Neutral | Up | Down |
| --- | --- | --- | --- | --- | --- | --- |
| K562 | 0.1 | 30 | 49108 | 48702 | 213 | 193 |
| K562 | 0.3 | 30 | 73383 | 72887 | 322 | 174 |
| K562 | 1 | 30 | 83653 | 82543 | 668 | 442 |
| K562 | 3 | 30 | 83665 | 80915 | 1770 | 980 |
| K562 | 10 | 30 | 83602 | 82381 | 567 | 654 |
| K562 | 30 | 30 | 83660 | 82311 | 552 | 797 |
| K562 | 100 | 30 | 83625 | 82191 | 505 | 929 |
| K562 | 300 | 30 | 39301 | 38737 | 323 | 241 |
| K562 | 1000 | 30 | 39329 | 38812 | 230 | 287 |
| K562 | 3000 | 30 | 10248 | 10086 | 97 | 65 |
| K562 | 10000 | 30 | 34540 | 33761 | 442 | 337 |
| MDA-MB-175 | 0 | 120 | 5395 | 5395 | non | non |
| MDA-MB-175 | 1 | 120 | 5392 | 5336 | 31 | 25 |
| MDA-MB-175 | 3 | 120 | 5393 | 5327 | 24 | 42 |
| MDA-MB-175 | 10 | 120 | 5392 | 5323 | 13 | 56 |
| MDA-MB-175 | 30 | 120 | 5392 | 5326 | 22 | 44 |
| MDA-MB-175 | 100 | 120 | 5394 | 5349 | 7 | 38 |
| MDA-MB-175 | 300 | 120 | 5390 | 5082 | 51 | 257 |
| MDA-MB-175 | 1000 | 120 | 5391 | 5117 | 36 | 238 |
| MDA-MB-175 | 3000 | 120 | 5390 | 4927 | 91 | 372 |
| MDA-MB-175 | 10000 | 120 | 5388 | 4724 | 193 | 471 |
| RPMI8226 | 0 | 60 | 24768 | 24768 | non | non |
| RPMI8226 | 0 | 120 | 23524 | 23524 | non | non |
| RPMI8226 | 0 | 240 | 20814 | 20814 | non | non |
| RPMI8226 | 0 | 480 | 17776 | 17776 | non | non |
| RPMI8226 | 0 | 960 | 26439 | 26439 | non | non |
| RPMI8226 | 0.1 | 60 | 24758 | 23782 | 286 | 690 |
| RPMI8226 | 0.1 | 120 | 23504 | 23091 | 145 | 268 |
| RPMI8226 | 0.1 | 240 | 20784 | 20629 | 88 | 67 |
| RPMI8226 | 0.1 | 480 | 17764 | 17577 | 136 | 51 |
| RPMI8226 | 0.1 | 960 | 26384 | 25873 | 285 | 226 |
| RPMI8226 | 1 | 60 | 24754 | 23732 | 72 | 950 |
| RPMI8226 | 1 | 120 | 23502 | 22969 | 137 | 396 |
| RPMI8226 | 1 | 240 | 20791 | 20653 | 90 | 48 |
| RPMI8226 | 1 | 480 | 17764 | 17483 | 247 | 34 |
| RPMI8226 | 1 | 960 | 26402 | 25902 | 344 | 156 |
| RPMI8226 | 10 | 60 | 24755 | 24164 | 42 | 549 |
| RPMI8226 | 10 | 120 | 23515 | 22862 | 242 | 411 |
| RPMI8226 | 10 | 240 | 20786 | 20407 | 276 | 103 |
| RPMI8226 | 10 | 480 | 17762 | 16989 | 681 | 92 |
| RPMI8226 | 10 | 960 | 26431 | 18988 | 4320 | 3123 |
| RPMI8226 | 100 | 60 | 24758 | 23638 | 63 | 1057 |
| RPMI8226 | 100 | 120 | 23513 | 22870 | 392 | 251 |
| RPMI8226 | 100 | 240 | 20800 | 19226 | 1227 | 347 |
| RPMI8226 | 100 | 480 | 17764 | 14219 | 2101 | 1444 |
| RPMI8226 | 100 | 960 | 26413 | 12412 | 6414 | 7587 |
| RPMI8226 | 1000 | 60 | 24761 | 23651 | 294 | 816 |
| RPMI8226 | 1000 | 120 | 23515 | 21892 | 896 | 727 |
| RPMI8226 | 1000 | 240 | 20802 | 18564 | 1666 | 572 |
| RPMI8226 | 1000 | 480 | 17766 | 13761 | 2321 | 1684 |
| RPMI8226 | 1000 | 960 | 26416 | 11652 | 6594 | 8170 |

#### Feature Representation

Protein PTM site representations are derived from ESM2 (37), a protein language model (PLM) that encodes residue-level sequence context. Chemical representations are obtained from UniMol (54), a pretrained molecular encoder that captures three-dimensional molecular geometry. For the machine learning (ML) models, the ESM2 and UniMol embeddings are concatenated into a single feature vector. For the deep learning (DL) model, the two modalities are processed by separate encoder branches each consisting of a series of multi-layer perceptron (MLP) layers and fused at a later stage for the final classification.

#### Models

We evaluate six model configurations spanning two families. In the **ML family**, we train a decision tree (DT) and a random forest (RF) on the full, unbalanced training set. We additionally train ensemble variants (DT-ensemble, RF-ensemble) in which the neutral class is partitioned into non-overlapping subsets, each combined with all minority-class examples to form a balanced training subset. In this setting, a separate model is trained on each subset and predictions are aggregated by majority vote. In the **DL family**, we train a dual-encoder model with standard cross-entropy loss (Dual Encoder) and with focal loss (36) (Dual Encoder Focal), which down-weights well-classified majority examples to focus learning on the minority classes.

#### Evaluation Metrics

The primary evaluation metrics are **Macro F1** and **Matthews Correlation Coefficient (MCC)**, which together capture both per-class balance and overall predictive quality under imbalance. Supporting metrics include per-class recall, precision, and area under the precision-recall curve (AUPRC) for the Up and Down classes, which are reported to characterize minority-class detection independently. Accuracy is not reported, as it is dominated by the neutral majority class and provides no meaningful signal in this imbalanced setting.

### Evaluation of Machine Learning and Deep Learning Models

#### Imbalance ratio, not model choice, is the dominant driver of performance

Across all three settings, the spread in Macro F1 among the six models within a single regime is at most 0.085 (Medium), whereas the shift in MCC between the Severe and Low regimes, holding the model family constant, is far larger in relative terms. In the Severe setting (K562, 0.1 *µ*g, ~100:1 ratio), all six models produce MCC values between −0.013 and 0.036, a range that is statistically indistinguishable from random. In the Low setting (RPMI8226, 100 *µ*g, ~5:1 ratio), MCC rises to 0.013–0.047 and minority-class recall reaches up to 0.43. This pattern indicates that the benchmark difficulty is primarily determined by the underlying biology, that is how many PTM sites are genuinely regulated at a given dose, rather than choice of classifier. Reducing the effective imbalance ratio, whether through higher drug doses, longer treatment windows, or targeted enrichment of regulated sites, may therefore yield larger gains than algorithmic improvements alone (3).

#### Rebalancing and focal loss trade precision for recall in opposite ways, and neither dominates

Ensemble rebalancing (DT-ensemble, RF-ensemble) and focal loss (Dual Encoder Focal) both shift the precision–recall trade-off toward higher recall, but through different mechanisms and with different failure modes. In the Severe setting, DT-ensemble raises Recall-Up from 0.000 to 0.235 but collapses Precision-Up to 0.005 (199 of 200 predicted Up events are false positives), while Dual Encoder Focal achieves zero recall on both minority classes and a negative MCC (−0.013). In the Low setting, the picture reverses: Dual Encoder Focal achieves the best MCC (0.047) and the highest combined recall (Recall-Up 0.431, Recall-Down 0.337), while RF-ensemble achieves the highest Macro F1 (0.334) but with Recall-Down of only 0.059. The absence of a single method that dominates across both metrics and all regimes is itself a key finding: it implies that the choice of intervention should be guided by the downstream use case, high-recall screening versus high-precision target prioritization, rather than by a single aggregate score (42).

#### The Dual Encoder detects Up-regulated sites more reliably than Down-regulated ones, except under focal loss

Averaged across models, Recall-Up exceeds Recall-Down in all three settings (Severe: 0.088 vs. 0.071; Medium: 0.147 vs. 0.108; Low: 0.298 vs. 0.175). This asymmetry is consistent with the biology of drug-induced phosphorylation: kinase activation events tend to produce large, reproducible fold-changes that are easier to distinguish from background noise, whereas phosphorylation loss can reflect kinase inhibition, phosphatase activation, or indirect network rewiring through distinct and more heterogeneous mechanisms (26). Focal loss partially reverses this asymmetry: in the Medium setting, Dual Encoder Focal achieves Recall-Down of 0.376 versus Recall-Up of 0.093, suggesting that the loss modulation preferentially rescues the Down class in this regime. Understanding why focal loss selectively amplifies Down-class detection, and whether this reflects a feature-space property of dephosphorylation sites, is an open question for future work.

#### A degenerate high-precision solution emerges in the Low setting, revealing a feature-space artifact

The standard RF in the Low setting achieves Precision-Up of 1.000 with Recall-Up of 0.001, meaning it predicts exactly one Up event in the entire test set and that prediction happens to be correct. This is not a useful classifier, but it is informative: it suggests that a small subset of Up-regulated sites has a feature signature that is highly separable from the Neutral class, while the vast majority of regulated sites are not. This pattern is consistent with the known structure of drug-induced PTM networks, where a small number of direct substrate sites (e.g., activation loop phosphorylations on the primary drug target) produce strong, reproducible signals, while the broader network of indirect regulation events is far more variable (27). Future models that explicitly distinguish direct from indirect regulation, for example by incorporating kinase-substrate network priors, may be able to exploit this separability more broadly.

## Conclusion

DrugPTM-Bench establishes the first integrated benchmark dataset that simultaneously advances drug discovery and confronts extreme data imbalance in machine learning. This resource addresses critical gaps in proteomics and pharmacology by systematically aggregating dose- and time-resolved post-translational modification (PTM) data across seven cell lines. To our knowledge, DrugPTM-Bench is the first dataset to provide dose- and time-resolved PTM regulation across multiple cell lines, offering a new paradigm for PTM-centric drug discovery. By capturing dynamic PTM regulation at the systems level, our dataset enables both biological discovery and computational modeling, facilitating a deeper understanding of drug-induced PTM dynamics and enhancing predictive modeling for therapeutic applications. Our evaluation demonstrates severe performance degradation in traditional methods (random forests, decision trees) and modern neural networks with focal loss when handling extreme class imbalance inherent to PTM regulation events. As a dual-purpose resource, DrugPTM-Bench promotes drug discovery and development through insights into chemical-perturbed cellular phenotypes and establishes a standardized framework for benchmarking next-generation imbalanced learning algorithms in biological systems.

## Data Availability

The DrugPTM-Bench dataset, benchmark splits, and associated metadata will be made publicly available upon acceptance of this manuscript.

## Acknowledgment

This project has been funded with federal funds from the National Institute of General Medical Sciences of the National Institute of Health (R01GM122845), the National Institute on Aging of the National Institute of Health (R01AG057555, R21AG083302), and the National Science Foundation (2226183).

## Additional Predictive Tasks Enabled by DrugPTM-Bench

While the primary benchmark task in this work is PTM regulation classification (T1), the multi-dimensional structure of DrugPTM-Bench. The benchmark encompasses pEC50 potency values, full dose-response curves, drug perturbation fingerprints across thousands of sites, and site-level sequence context enabling a family of additional drug discovery relevant tasks. These tasks are formally defined here to establish a community roadmap and invite future contributions. They are not benchmarked in the present work. Each task fills a gap not addressed by existing benchmarks.

### T2: pEC50 Potency Regression

In decryptM (51), each PTM site × drug combination with a measurable dose-response is fitted across 11 drug concentrations to yield a pEC50 value. The negative log_10_ of the molar concentration at which the signal ratio reaches half its maximum effect. The T2 task is to predict this pEC50 from protein sequence context, drug structure, cell line identity, bypassing the need to run the full 11-point dose-response assay. This is fundamentally different from the canonical drug-target affinity benchmarks Davis and KIBA (47), which predict thermodynamic binding affinity (K_d_ or K_i_) measured in cell-free biochemical assays. The pEC50 in DrugPTM-Bench is a *functional cellular potency* : it reflects not only direct binding but also downstream pathway buffering, feedback regulation, and cell-line-specific signaling wiring (51). A drug can have high binding affinity for a kinase yet low pEC50 on a downstream phosphosite if the pathway is buffered by compensatory mechanisms, a distinction that cell-free affinity measurements cannot capture. From a drug discovery perspective, a model that accurately predicts pEC50 at PTM-site resolution could prioritize which drug-site pairs to validate experimentally, reducing the cost of dose-response phosphoproteomics by an order of magnitude. An important limitation is that pEC50 values are only defined for regulated sites (~47,000 site × drug pairs in DrugPTM-Bench), making the training set smaller than Davis or KIBA, smaller data regimes are very common in computational drug discovery.

### T3: Mechanism-of-Action Prediction from PTM Profile

Drugs with the same mechanism of action perturb overlapping sets of PTM sites: kinase inhibitors suppress phosphorylation of their direct substrates and downstream effectors, HDAC inhibitors broadly alter histone and non-histone acetylation, and proteasome inhibitors trigger widespread phosphorylation changes through proteostatic stress (51; 16). A PTM perturbation fingerprint, the vector of signal ratios across all detected sites for a given drug in a given cell line, is therefore mechanistically closer to drug action than transcriptomic signatures, because PTMs are the direct molecular readout of kinase, phosphatase, acetyltransferase, and deacetylase activity rather than a downstream transcriptional consequence. The T3 task is to predict a drug’s mechanism class from its PTM fingerprint. Existing MoA prediction benchmarks, including LINCS L1000 (46), MoAble (23), MOASL (24), and the PANACEA DREAM Challenge (13), all use transcriptomic perturbation signatures as input, and none use PTM-level profiles. Mitchell et al. (41) profiled 875 compounds by quantitative proteomics (protein expression, not PTM) and demonstrated MoA clustering, but PTM resolution was not exploited. With 27 drugs spanning eight mechanism classes in DrugPTM-Bench, T3 is a tractable but non-trivial classification problem that could reveal whether PTM fingerprints encode MoA information not captured by transcriptomics.

### T4: Drug-Specific PTM Site Sensitivity Ranking

Mass spectrometry-based phosphoproteomics experiments routinely detect thousands of PTM sites per condition, but experimental follow-up (mutagenesis, functional assays, structural studies) can only be performed on a handful of sites (42). A computational model that ranks PTM sites by their predicted sensitivity to a specific drug would directly reduce this experimental bottleneck by prioritizing the most drug-responsive sites for validation. Existing PTM prioritization methods rank sites by intrinsic functional importance: Beltrão et al. (3) used evolutionary conservation and structural context. Ochoa et al. (42) integrated 6,801 proteomics experiments to build a functional landscape of the human phosphoproteome. Lastly, Kennedy et al. (27) developed PTM-centric base editor screens to assess phosphosite functionality at scale experimentally. All of these approaches are drug-agnostic, they rank sites by their general biological importance, not by their sensitivity to a particular perturbation. DrugPTM-Bench enables the first drug-specific PTM site sensitivity ranking benchmark, where the input is a drug fingerprint paired with a protein sequence and all its candidate PTM site contexts, and the output is a ranked list of sites by predicted |signal ratio|. The Johnson et al. kinome substrate specificity atlas (26), which maps the sequence preferences of over 300 human Ser/Thr kinases, provides a natural source of sequence-level priors for T4 models targeting phosphorylation sites. Appropriate evaluation metrics for T4 include normalized discounted cumulative gain (NDCG) and Spearman correlation between predicted and observed |signal ratio| rankings per drug-protein pair.

## PTM Event Regulation Distribution (T1 data distribution)

